# The phage shock protein response of *Listeria monocytogenes* influences tolerance to the multipeptide bacteriocin garvicin KS

**DOI:** 10.1101/2025.10.08.681093

**Authors:** Thomas F. Oftedal, Trond Løvdal, Morten Kjos

## Abstract

An estimated 30% of all food produced worldwide is lost or wasted every year. A considerable portion of that waste is due to perishable food products, that spoil and can become unsafe to eat relatively quickly. For some high-quality perishables, such as fresh fish and cold-smoked salmon, traditional food preservation techniques are unsuitable as they can compromise sensory qualities such as flavor, texture, and freshness. These products often support the growth of the human pathogen *Listeria monocytogenes*, which can be present if thermal treatment is not applied. Thus, antilisterial bacteriocins like garvicin KS (GarKS) in combination with conventional technologies like high-pressure processing (HPP) or modified atmosphere packaging (MAP) are being investigated as hurdle strategies to increase the shelf life and food safety of packaged fish products. In this study we showed that *L. monocytogenes* strains associated with fish food and fish processing plants are susceptible to GarKS with MIC values ranging from 20 to 275 nM. Exposure to GarKS resulted in an upregulation of genes involved in the phage shock protein response, and isolation of resistant mutants indicated a low frequency of resistance to GarKS (10□□ to 10□^11^). Resistant mutants were shown to harbor disruption mutations in *lmo2468*, encoding a PspC-domain-containing protein. Overexpression of this gene increased susceptibility to GarKS two-fold and restored wild-type susceptibility in a disruption mutant. This study identify the phage shock protein response as a key player involved in susceptibility to GarKS.

## Introduction

An estimated 1.05 billion metric tons of food was wasted in 2022, a number that is expected to increase in the coming years (Gatto and Chepeliev, 2024; United Nations Environment Programme, 2024). In the EU, over half of all food waste occurs in households, and about 25-50% of household food waste is attributed to suboptimal food packaging (Silvenius et al., 2014; Uhlig et al., 2025; Williams et al., 2020). Fresh products such as meat and fish are particularly susceptible to sensory decline and spoilage. However, waste of these products can be reduced by using packaging methods that inhibit the growth of spoilage microorganisms and foodborne pathogens or protect them from contamination (Verghese et al., 2015).

Some fish products, such as cold-smoked salmon, have a recommended shelf life of only a few days when opened, even at refrigeration temperatures, due to the potential of *Listeria monocytogenes* contamination (Huang et al., 2023). Fish and seafood products, including smoked fish, are one of the most frequent sources of human listeriosis and are therefore highly relevant to public health (Koutsoumanis et al., 2022). *L. monocytogenes* is widespread in nature and frequently found on animals, in soil, manure, water, and food processing plants (Kathariou, 2002; Stea et al., 2015; Vivant et al., 2013; Zakrzewski et al., 2024). The organism is very tolerant to environmental stress such as salt content and is able to grow at refrigeration temperatures (Chan and and Wiedmann, 2008; Ribeiro and Destro, 2014). Furthermore, *L. monocytogenes* is able to form biofilms on various surfaces and persist in food processing plants despite rigorous cleaning routines (Mędrala et al., 2003; Zakrzewski et al., 2024). Consequently, there is a need for food preservation techniques that can eliminate *L. monocytogenes* from fresh and ready-to-eat foods, particularly fish products.

There has been an increasing interest in novel food preservation techniques, including biopreservation with natural antimicrobials, such as bacteriocins, and high-pressure processing (HPP). Bacteriocins are bacterially produced antimicrobial peptides that may or may not be post-translationally modified, of which many are active against *L. monocytogenes*. In particular, the so-called pediocin-like class IIa bacteriocins are known for their extremely potent antilisterial activity (Bédard et al., 2018; Katla et al., 2003). HPP is a preservation technique that typically involves exposure to a high hydrostatic pressure of typically 100 to 600 MPa for one to about five minutes, which results in the inactivation of microorganisms such as *L. monocytogenes* (Jiménez-Colmenero and Borderias, 2003; Wiśniewski et al., 2024).

HPP treatment of fish can increase hardness and chewiness and lighten the appearance of the meat (Zhao et al., 2019). These unwanted changes become more pronounced at higher pressures and treatment times (Zhao et al., 2019). Thus, *L. monocytogenes* is not completely eliminated during HPP treatments commonly used as part of food processing (Lakshmanan and Dalgaard, 2004; Wiśniewski et al., 2024). Certain strains of *L. monocytogenes* have been shown to be more tolerant to pressure treatment (barotolerant) than others, for example, *L. monocytogenes* RO15 showed no significant reduction after treatment at 400 MPa (Duru et al., 2020). One idea is therefore to combine bacteriocins with HPP to further eliminate bacteria from packaged foods and allow for faster treatments and lower pressure (Løvdal, 2015). The bacteriocin pediocin PA-1 has been shown to act synergistically with HPP treatment in reducing *L. monocytogenes* (Komora et al., 2020). However, for this bacteriocin, frequent resistance development is a major issue (Gravesen et al., 2002a).

Bacteria may become resistant to a bacteriocin by spontaneous mutations that disrupt or alter a protein that is required by the bacteriocin to exert its antimicrobial potential. Bacteriocins such as pediocin PA-1, which is commercialized under the brand name ALTA 2341 as a food preservative by the Kerry Group plc (Ireland), kill by binding to the mannose phosphotransferase system (Man-PTS)—a sugar transporter that is ubiquitous in bacteria (Kjos et al., 2009; Tymoszewska and Aleksandrzak-Piekarczyk, 2024). Mutations in the gene encoding a subunit of Man-PTS can render cells non-susceptible to bacteriocins targeting Man-PTS (Kjos et al., 2009; Oftedal et al., 2021; Tymoszewska and Aleksandrzak-Piekarczyk, 2024). In contrast, nisin, the only bacteriocin widely used as a food preservative today (Demirgül et al., 2025), binds to a non-proteinaceous molecule, lipid II, to exert its antimicrobial activity (Brötz et al., 1998). Despite this, resistance to nisin in *L. monocytogenes* can arise by spontaneous mutations and occurs at a frequency from 10^−6^ to 10^−3^ in strains isolated from cheese (Martínez et al., 2005). The widely used strain *L. monocytogenes* Scott A develops nisin resistance at frequencies from 10^−6^ to 10^−8^ when subjected to nisin at 2-to 8 times the minimal inhibitory concentration (MIC) (Ming and Daeschel, 1993). Nisin resistance in *L. monocytogenes* is a complex phenotype that involves cell envelope stress response systems, cell envelope modifying proteins, and cell wall biosynthesis proteins (Bergholz et al., 2013; Gravesen et al., 2004).

Garvicin KS (GarKS) is a bacteriocin produced by *Lactococcus garvieae* KS1546 belonging to a class of unmodified and leaderless bacteriocins (Ovchinnikov et al., 2016). GarKS consists of three different peptides (GakABC) that must be present in equal amounts for optimal activity (Ovchinnikov et al., 2016). Unmodified and leaderless bacteriocins are in their active form immediately after translation. GarKS shows potent antimicrobial activity against the food pathogenic species *S. aureus* and *L. monocytogenes* with MIC values as low as 160 nM for some strains (Ovchinnikov et al., 2016). One of the major challenges to the use of bacteriocins for food preservation is production costs and low yields (Liang et al., 2025). The success of nisin as a commercial product is arguably in large part due to the high production potential of the producer strain using inexpensive milk-based growth medium (de Arauz et al., 2009; Zhao et al., 2021; Zhou et al., 2008). Nisin is produced commercially by batch fermentation of milk, which yields a fermentate containing 2.5% (wt/wt) nisin (Demirgül et al., 2025). Similar high-yield production has also been demonstrated for GarKS using milk-based growth media (Telke et al., 2019), and this bacteriocin is therefore considered particularly interesting for development as a biopreservative. However, a major concern for antimicrobials used for industrial, agricultural, and medical purposes is the development of resistance.

The molecular target exploited by GarKS is not known, and most leaderless bacteriocins are thought to not require a receptor for killing and resistance to GarKS in the foodborne pathogen *L. monocytogenes* have not been studied (Ovchinnikov et al., 2020; Perez et al., 2018). The aim of this study was to assess the susceptibility of *L. monocytogenes* to GarKS, with a particular emphasis on barotolerant strains and strains associated with fish food and fish processing plants. We estimate the frequency of resistance and identify potential resistance mechanisms to GarKS in the barotolerant *L. monocytogenes* RO15. Genes involved in GarKS susceptibility were identified by RNA sequencing and differential gene expression analysis. The direct involvement of candidate genes was confirmed by molecular cloning, gene expression, and isogenic complementation.

## Materials and methods

### Strains and growth conditions

*L. monocytogenes* was cultivated in Brain Heart Infusion broth (BHI; VWR International) at 37 °C with shaking (180 rpm). *E. coli* was grown in LB broth (10 g/L tryptone [Oxoid], 10 g/L NaCl [VWR], 5 g/L yeast extract [Oxoid]). Culture media for strains harboring pNZ-derived plasmids were supplemented with either 10 µg/ml chloramphenicol for *L. monocytogenes* or 20 µg/ml chloramphenicol for *E. coli*. All strains and plasmids used in this study are listed in Table 1.

**Table 1.**
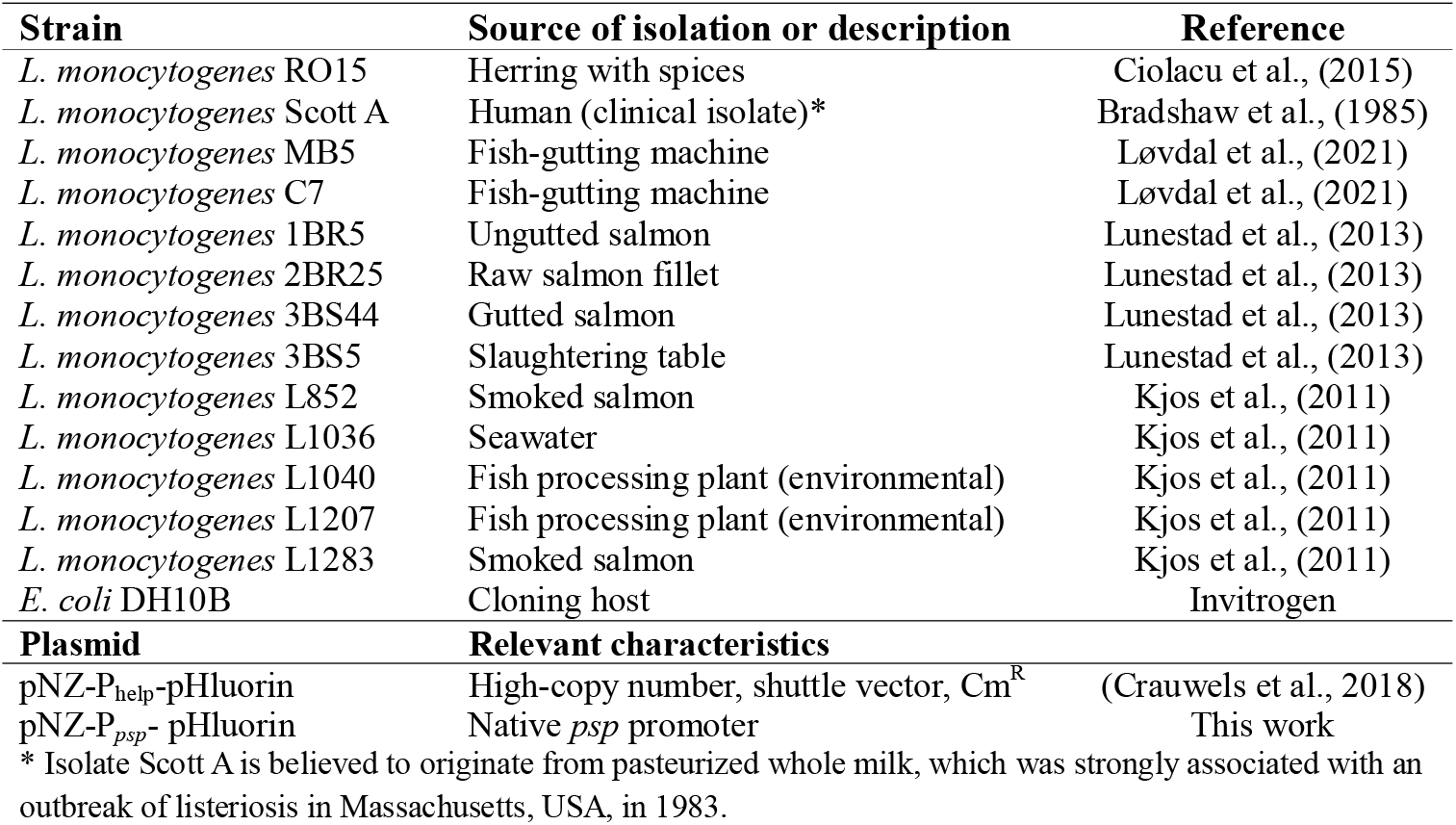
Strains and plasmids used in this study.

### Preparation of GarKS

The individual peptides comprising GarKS were purchased from PepMic Co., Ltd. (China) at a purity of >90%; each peptide was solubilized to a stock concentration of 3 mM using 0.1% TFA. All three peptides were combined in equal volumes to make a 1 mM solution of GarKS. Peptides were incubated at 50 °C and 1200 rpm using a Thermo-Shaker TS-100 (Biosan, Latvia) for 10 min prior to mixing.

### Minimum inhibitory concentration

Cultures of *L. monocytogenes* were grown for 16 h and then diluted 100-fold in the wells of a 96-well plate (Sarstedt, Germany) with BHI containing various concentrations of GarKS (serial dilution) to a total volume of 0.2 ml per well. The plate was incubated statically at 37 °C for 24 h before growth was measured using a SPECTROstar Nano (BMG Labtech, Germany) spectrophotometer at 600 nm. The MIC was defined as the lowest concentration of GarKS needed to inhibit growth by at least 90% compared to a control with no added antimicrobial. The MIC was determined as the mean of four independent assays.

### RNA isolation and sequencing

An overnight culture of *L. monocytogenes* RO15 was diluted 100-fold in brain heart infusion broth (BHI, VWR) and incubated at 37 °C with shaking (180 rpm) to an optical density at 600 nm of 0.5. The culture was then divided in two; garvicin KS (in 0.1% TFA) was added to one culture to a final concentration of 12.5 nM, while an equal volume of 0.1% TFA was added to the other (control). Cultures were then incubated for another 20 or 60 min before harvesting by centrifugation (4000g, 4 °C, 10 min). Supernatant was removed using a pipette, and the cell pellet was flash frozen in liquid nitrogen and stored at -80 °C for approximately 24 hours. RNA was isolated using a combination of the TRIzol Max Bacterial RNA Isolation Kit (Invitrogen) and the RNeasy Mini Kit (Qiagen). Firstly, frozen cells were thawed by resuspension in room temperature TRIzol reagent with 0.2 volumes of chloroform and transferred to FastPrep tubes containing 500 µl of zirconia beads. Cells were subjected to three rounds of bead beating in a FastPrep-24 (MP Biomedicals) at 6.5 m/s for 20 seconds with 1 min on ice between runs.

The resulting lysate was separated into an aqueous and organic phase by centrifugation (16000g, 15 min, 4 °C) and the upper aqueous phase was transferred to RNeasy spin columns. Bound RNA was washed according to the manufacturer’s instructions: once in buffer RW1 and twice in buffer RPE (8000 g, 15 s). Purified RNA was eluted with 50 µl of DEPC-treated Milli-Q water. Ribosomal RNA removal, stranded library preparation, and sequencing were performed by Novogene Ltd. (Cambridge, UK).

### Differential gene expression analysis

Reads were processed by fastp using the default settings. The genome sequence for *L. monocytogenes* RO15 was obtained from GenBank (Accession CADEHJ000000000.1). Reads were mapped to the reference genome using Bowtie2 with the very-sensitive option. Reads mapping to features was counted using htseq-count with the reverse stranded option (--stranded=reverse). Differential gene expression analysis was performed in R using DESeq2; statistical significance was based on the Benjamini-Hochberg adjusted p-values using an α of 5% (<0.05) as a threshold. After analysis, genes were renamed to the closest ortholog from *L. monocytogenes* EGD-e (accession AL591824) assisted by the OrthoFinder v2.5.5 software (Emms and Kelly, 2019).

### Isolation of GarKS-tolerant mutants

To isolate cells with increased tolerance to GarKS, 0.1 ml of a stationary phase culture of *L. monocytogenes* RO15 was directly plated on BHI agar containing GarKS at 2X, 4X, 8X, 10X, and 12X the MIC_90_ (corresponding to 250, 500, 1000, 1250, and 1500 nM). Plates were incubated overnight at 37 °C. To determine the frequency of tolerant cells, an equal aliquot of the same culture was serially diluted (10-fold dilutions), and 0.1 ml of the dilutions 10^−6^, 10-7, and 10^−8^ were plated. The dilution with 30 to 200 colonies was counted. The frequency of tolerant cells was calculated as the ratio of the number of colonies on plates containing GarKS over the number of cells (CFU) plated. The genomic region containing the *psp* locus was amplified by PCR using the primer pair Lm_psp_F and Lm_psp_R. Amplicons were sent to Eurofins Genomics for sequencing using the ONT lite clonal amplicon sequencing service. Sequence variants were further confirmed by Sanger sequencing (Eurofins Genomics) using the Lm_psp and Lm_psp_In primers listed in Table 2.

**Table 2.**
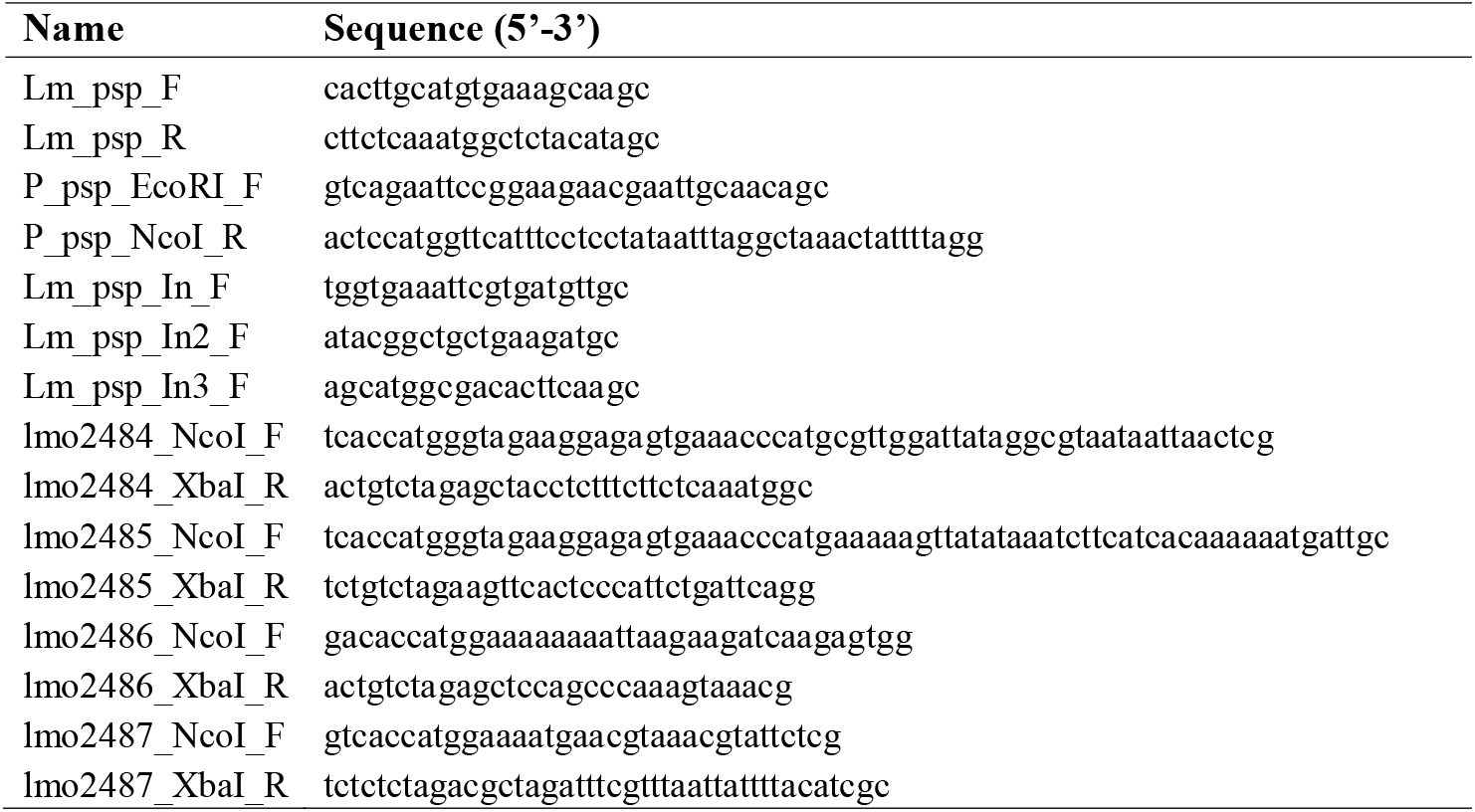
List of oligonucleotides used in this study.

### Preparation of electrocompetent *L. monocytogenes*

*L. monocytogenes* RO15 was made electrocompetent following the protocol by Monk et al. (2008) with some modifications. The strain was grown in BHI at 37 °C with shaking (180 rpm) for 16-20 h, then diluted 100-fold in 500 ml of BHI containing 500 mM sucrose. Incubation was continued until the culture reached an optical density at 600 nm of 0.3, measured using a 10-mm cuvette on a SPECTROstar Nano. Ampicillin was added to the culture to 20 µg/ml, and the culture was incubated for another 2 h. Growth was stopped by swirling the culture flask in an ice bath for 10 min. Cells were washed three times in ice-cold 10% glycerol with 500 mM sucrose by centrifugation (4000 g, 10 min, 4 °C), with a reduction in volume for each wash (500, 200, 50 ml). Cells were incubated at 37 °C for 20 min with 10 µg/ml lysozyme, then washed twice as described previously, first in 20 ml and then in 2 ml of 10% glycerol with 500 mM sucrose. Aliquots of 50 µl were frozen at -80 [°C until use.

### Construction of pNZ-P_psp_-pHluorin

The plasmid pNZ-P_help_-pHluorin was digested with NcoI and EcoRI (FastDigest, Thermo Scientific) to excise the P_help_ promoter region from the plasmid. The linearized vector was separated by gel electrophoresis and purified using the NucleoSpin Gel and PCR Clean-up Kit (Macherey-Nagel). The upstream region of the *psp* locus in *L. monocytogenes* RO15 (corresponding to *lmo2487*-*2484* in *L. monocytogenes* EGD-e, accession AL591824), containing the native *psp* promoter (P_*psp*_), was amplified using the primer pair P_psp_EcoRI_F and P_psp_NcoI_R and subsequently purified from the PCR reaction mixture with the NucleoSpin Gel and PCR Clean-up Kit (Macherey-Nagel).

The linearized vector and amplicon were combined in a 1:3 molar ratio with 1 U/µl T4 DNA ligase (Thermo Scientific) and the provided reaction buffer to a reaction volume of 30 µl. Ligation was performed for 16 h at 4 °C before inactivation by heating at 65 °C for 10 min. An aliquot of *E. coli* MAX Efficiency DH10B chemically competent cells (Invitrogen) were thawed on ice for 5 min, then mixed with 3 µl of the ligation mixture. The mixture was incubated on ice for 30 min, then placed in a water bath at 42 °C for 30 seconds, then back on ice for 5 min. LB medium was added to the cells and incubated for 1 h at 37 °C with shaking (180 rpm). Cells were plated on LB agar containing 10 µg/ml chloramphenicol for selection. The construct was sequenced by Eurofins Genomics for verification.

### Construction of *psp* overexpressing strains

Each gene of the *psp* locus (locus tags *OCPFDLNE_02658-02661* in *L. monocytogenes* RO15, or *lmo2484*-*2487* in *L. monocytogenes* EGD-e with accession AL591824) was amplified using the corresponding primer pairs listed in Table 2. Amplicons were purified from PCR reactions using the NucleoSpin Gel and PCR Clean-up Kit (Macherey-Nagel). The plasmid pNZ-P_psp_-pHluorin (described above) was digested with FastDigest XbaI and NcoI (Thermo Scientific) to excise the pHluorin gene. The plasmid backbone was separated by gel electrophoresis and purified similarly. Ligation of each gene into the linearized pNZ-P_psp_ vector and transformation into *E. coli* DH10B was performed as explained for the construction of pNZ-P_psp_-pHluorin above.

## Results

### *L. monocytogenes* from food-associated environments are sensitive to GarKS

The bacteriocin GarKS has previously been shown to inhibit two strains of *L. monocytogenes* (Ovchinnikov et al., 2016). To systematically assess the susceptibility of *L. monocytogenes* to GarKS, in particular isolates originating from food, fish, and fish processing environments, isolates were acquired from Nofima (Norwegian Institute of Food, Fisheries and Aquaculture Research) and our laboratory collection (Table 1). The isolates were then assessed for GarKS susceptibility using a minimum inhibitory concentration (MIC) assay. As shown in Table 3, the MIC for all strains tested (n=13) was between 20 nM for the most susceptible strain, L1207, and 275 nM for the least susceptible strain, C7, a difference of 14-fold. The strain RO15 showed an average susceptibility of 125 nM, while the mean MIC_90_ across all strains was 128 nM.

**Table 3.** Susceptibility of *L. monocytogenes* strains isolated from food, fish, and fish processing plants to GarKS.

| Strain | MIC<sub>90</sub> (nM) |
| --- | --- |
| *L. monocytogenes* RO15 | 125 |
| *L. monocytogenes* Scott A | 200 |
| *L. monocytogenes* MB5 | 225 |
| *L. monocytogenes* C7 | 275 |
| *L. monocytogenes* 1BR5 | 100 |
| *L. monocytogenes* 2BR25 | 150 |
| *L. monocytogenes* 3BS44 | 125 |
| *L. monocytogenes* 3BS5 | 125 |
| *L. monocytogenes* L852 | 80 |
| *L. monocytogenes* L1036 | 80 |
| *L. monocytogenes* L1040 | 80 |
| *L. monocytogenes* L1207 | 20 |
| *L. monocytogenes* L1283 | 80 |

### Resistance to GarKS is infrequent in *L. monocytogenes*

We further sought to investigate to what extent *L. monocytogenes* could develop resistance to GarKS. To do this, *L. monocytogenes* RO15 cultures were plated on agar containing various concentrations of GarKS (from 2-to 12 times the MIC_90_). Notably, resistance development was rare, and growth was only observed on plates containing 2-, 4, and 8 times the MIC_90_. The frequency (proportion or ratio) of cells able to grow on GarKS at these concentrations was in the range of 10^−9^ – 10^−11^ (Table 4). The highest concentration of GarKS permitting growth was 1000 nM (8X MIC_90_).

**Table 4.** Frequency of resistance development to various concentrations of GarKS expressed as the ratio of CFU growing at the stated GarKS concentration over the total CFU plated.

| GarKS (nM) | Ratio of total cells |
| --- | --- |
| 250 | $1.5 \cdot 10^{-9}$ |
| 500 | $1.3 \cdot 10^{-10}$ |
| 1000 | $1.6 \cdot 10^{-11}$ |

In total, 17 isolated resistant colonies, including one from 1000 nM, 11 from 500 nM, and five from 250 nM, were reassessed for GarKS susceptibility using the MIC assay and compared to the wild-type MIC_90_ (125 nM). Surprisingly, six of the isolates showed the same susceptibility as the wild-type strain, and seven isolates were slightly more tolerant with an MIC_90_ of 200 nM (1.6-fold higher than the wild type; mean of four independent assays). The remaining four isolates had an MIC_90_ of 250 nM, despite all being isolated from plates containing 1000 nM GarKS.

### The Phage Shock Protein response is activated by GarKS

Given the low frequency and level of GarKS resistance development observed above, the potential for resistance development was also investigated by transcriptomics. RNA was isolated from mid-log phase cultures of the barotolerant strain *L. monocytogenes* RO15 after exposure to 0.1X MIC_90_ for 20 min. *L. monocytogenes* have been shown to alter transcription of genes in 5-15 minutes (Arvaniti et al., 2025; Nikparvar et al., 2021; Piveteau et al., 2011). The genes most significantly upregulated by GarKS exposure are shown in Figure 1. These 12 genes showed a log_2_ fold change higher than 1 and a p-value less than 0.05.

**Figure 1.**
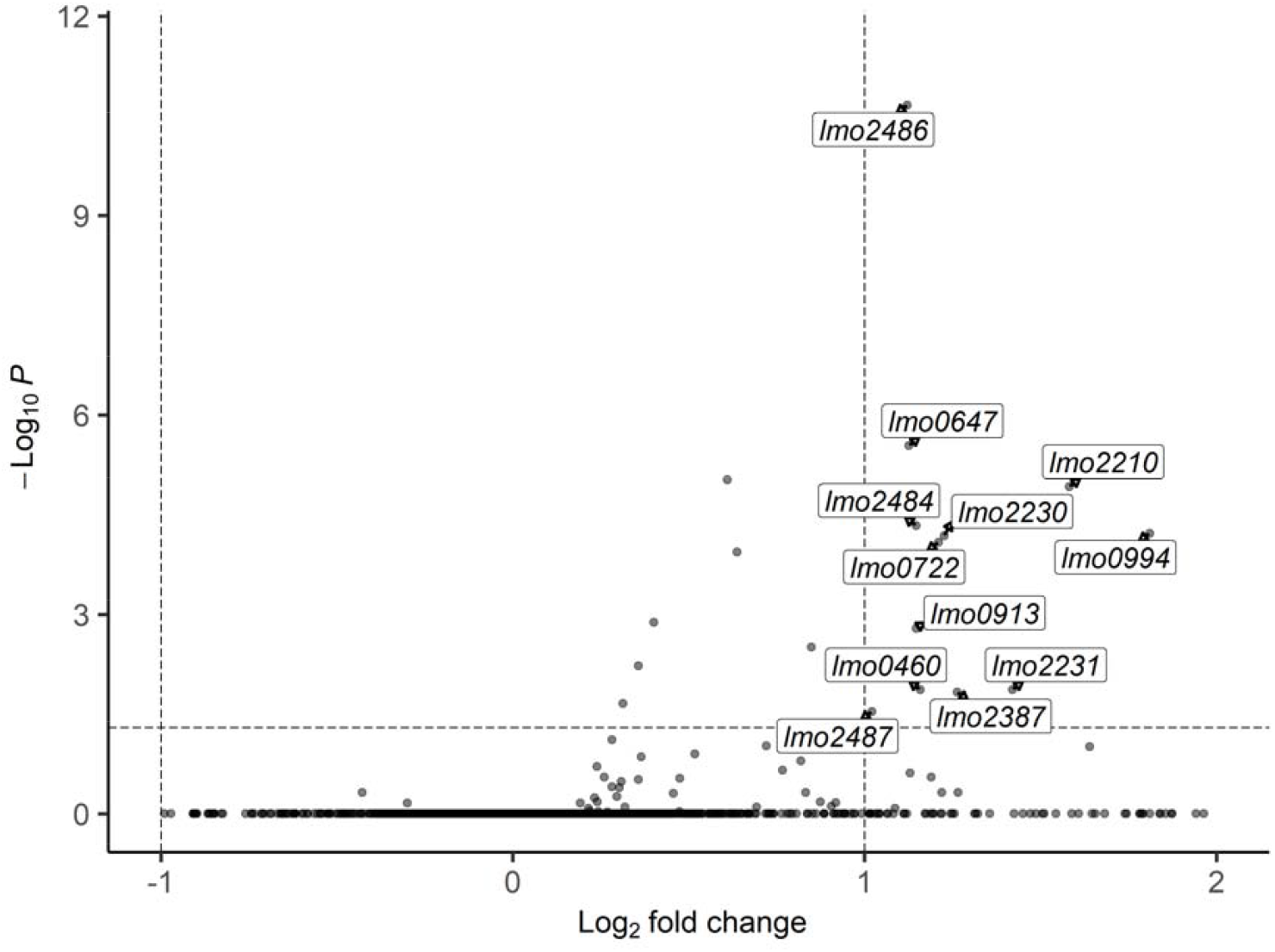
Volcano plot of differentially expressed genes from GarKS exposure in *L. monocytogenes* RO15 after 20 min. Genes in the upper right quadrant have a p-value less than 0.05 and a log2 fold change of more than 1. The figure was generated using the EnhancedVolcano package for R (Blighe et al., 2019).

In total, GarKS exposure resulted in the upregulation of 22 genes (Table 5) putatively belonging to 17 operons. The gene most significantly differentially expressed was *lmo2486*, encoding a “toastrack”-domain and phage shock protein C (PspC)-domain-containing protein. Two other genes, *lmo2484* (encoding a phage holin domain-containing protein) and *lmo2487* (encoding another toastrack-domain-containing protein), predicted to be in the same operon, were also upregulated upon GarKS exposure.

**Table 5.**
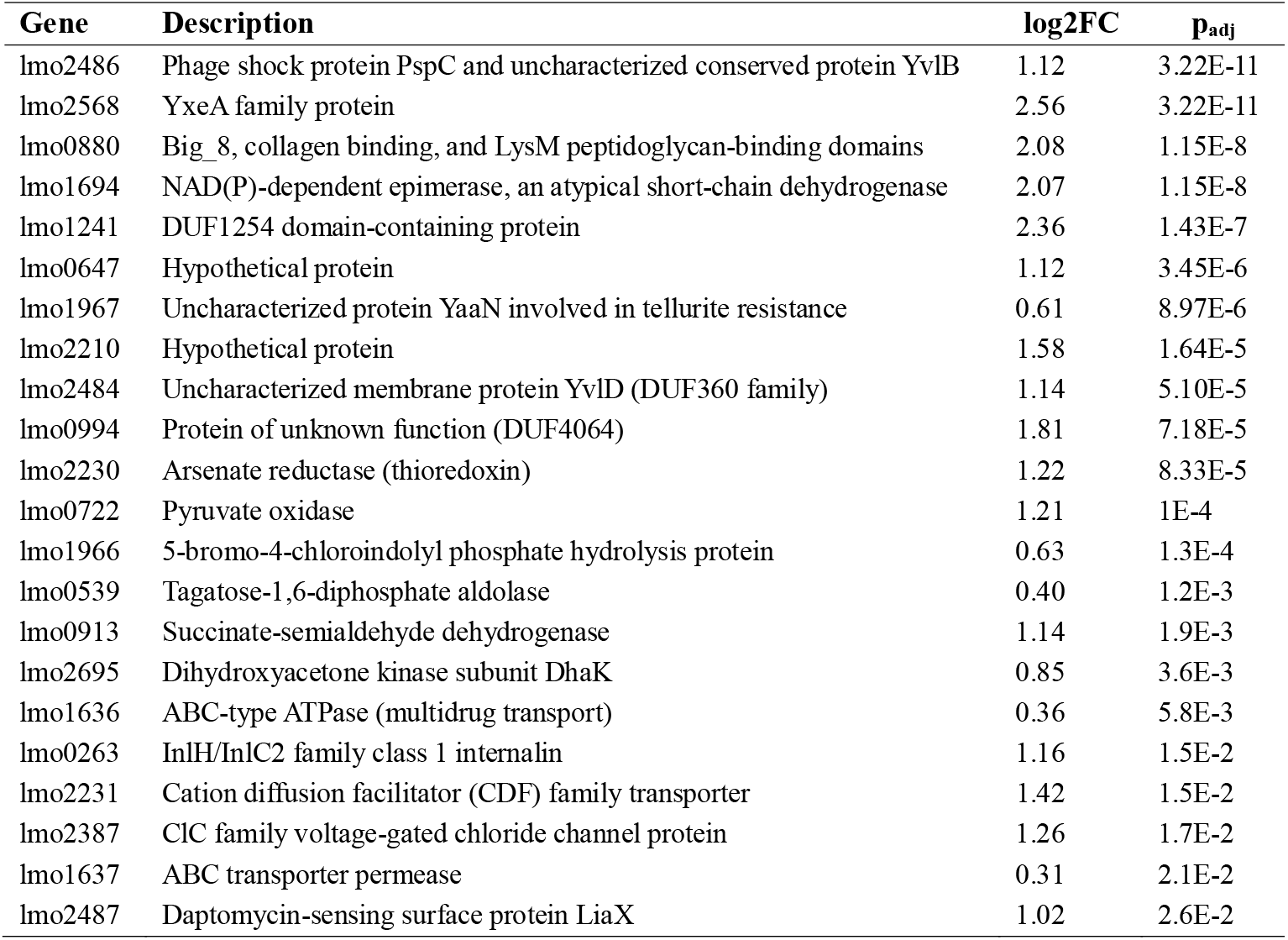
Differentially expressed genes (adjusted p-value < 0.05) found in *L. monocytogenes* RO15 after GarKS exposure (12.5 nM) for 20 min. Genes were named using the locus tags from *L. monocytogenes* EGD-e (Accession no. AL591824).

**Table 6.** Susceptibility to GarKS of *L. monocytogenes* RO15 expressing the Psp proteins from a plasmid and an *lmo2486* disruption mutant (GM1) expressing *lmo2486* (complemented). A plasmid expressing a pH-sensitive variant of green fluorescent protein (pNZ-P_psp_-pHluorin) was used as a control.

| Strain | MIC<sub>90</sub> (nM) |
| --- | --- |
| *Lmo* RO15 pNZ-P<sub>psp</sub>-*lmo2487* | 80 |
| *Lmo* RO15 pNZ-P<sub>psp</sub>-*lmo2486* | 40 |
| *Lmo* RO15 pNZ-P<sub>psp</sub>-*lmo2485* | 80 |
| *Lmo* RO15 pNZ-P<sub>psp</sub>-*lmo2484* | 80 |
| *Lmo* RO15 control | 80 |
| *Lmo* GM1 control | 250 |
| *Lmo* GM1 pNZ-P<sub>psp</sub>-*lmo2486* | 80 |

Other operons that were found to be upregulated were an ABC transporter, *lmo1636-1637*, and cation efflux systems, *lmo1966-1967* and *lmo2230-2231*. The most overexpressed gene (highest fold-change difference) was *lmo2568*, a YxeA family protein.

### GarKS-tolerant mutants harbor mutations in a PspC-domain containing protein

Given that the *lmo2487*-*6-5-4* operon was shown to be upregulated, this region of the genome was amplified and sequenced in the four isolates exhibiting the highest increase in GarKS tolerance. Strikingly, all four isolates were found to have a nonsense mutation in the gene encoding Lmo2486. All four mutants had the same deletion of an adenine in the 12th position in the gene (c.12delA), despite the fact that the four resistant mutants were isolated in separate experiments. The consequence of the deletion is a frameshift that introduces a stop in the 5th codon (p.K4NfsX).

### Lmo2486 is involved in GarKS susceptibility

To further investigate the involvement of Lmo2486 in GarKS susceptibility, the gene was cloned on a high-copy number plasmid pNZ-P_help_ and expressed in *L. monocytogenes* RO15 wild type and one of the *lmo2486* frameshift mutants mentioned above (mutant hereafter designated GM1). Initially, the genes were placed behind a strong constitutive promoter, P_help_ (highly expressed *Listeria* promoter), but no transformants could be obtained. As such, the promoter was replaced by the native promoter (P_psp_) of the *psp* operon, which allowed us to obtain correct transformants. Complementing the resistant Lmo2486 disruption mutant GM1 with the intact gene on the pNZ-P_psp_ plasmid caused the strain to become more susceptible to GarKS, with the MIC_90_ decreasing from 250 nM to 80 nM (3-fold decrease). Furthermore, *L. monocytogenes* RO15 harboring pNZ-P_psp_-*lmo2486* showed a 2-fold reduction in MIC_90_ (mean of 6 independent assays) compared to a control strain, showing that the protein influences tolerance to GarKS.

Furthermore, we also expressed the other proteins in the *psp* operon (Lmo2487, Lmo2485, Lmo2484) in the same manner; however, this did not affect susceptibility compared to the control.

## Discussion

In this study we examined the GarKS susceptibility of strains associated with fish food and processing plants. We further quantified resistance development in a barotolerant strain, *L. monocytogenes* RO15, which might be a more representative strain for surviving cells following HPP treatment.

The susceptibility of strains associated with fish food and processing plants to GarKS was between 20 and 275 nM. In comparison, the reported MIC of nisin in various strains of *L. monocytogenes* is around 2 µM, suggesting a potency of GarKS which is from 7-to 100 times higher than that of nisin (Martínez and Rodríguez, 2005). As a comparison with antibiotics, concentrations needed to inhibit most strains of *L. monocytogenes* isolated from fish and water sources vary from 10 to 20 µM (Rodas-Suárez et al., 2006). Although some bacteriocins like pediocin PA-1 exhibit even higher potency than GarKS with MIC values typically below 10 nM, however, this bacteriocin has the problem of chemical instability and rapid resistance development (from 10^−4^ to 10^−6^) (Bédard et al., 2018; Gravesen et al., 2002b). On the other hand, we here show that *L. monocytogenes* does not develop resistance against GarKS rapidly, with resistance frequencies less than 10^−9^.

Furthermore, the level of resistance for the isolated colonies was low; at maximum an 8-fold increase in MIC_90_ was observed despite repeated efforts to isolate mutants with higher resistance levels. We could not obtain any cells of *L. monocytogenes* RO15 that could grow at GarKS concentrations above 1000 nM (8X MIC). The inability to isolate mutants with a higher resistance to GarKS supports the preconception that most leaderless bacteriocins, such as GarKS, do not target a specific protein on the target cell for killing (Perez et al., 2018). The only known exception is the LsbB family of leaderless bacteriocins, which is known to target a zinc metalloprotease (RseP/RasP/YvjB) on target cells for killing (Kristensen et al., 2022; Ovchinnikov et al., 2017; Uzelac et al., 2013). However, a molecular target of GarKS that is not a protein cannot be excluded, as for example nisin which kills cells via an interaction with lipid II, or cinnamycin which interacts with phosphatidylethanolamine (Brötz et al., 1998; Machaidze and Seelig, 2003).

All four isolates with the highest increase in tolerance to GarKS showed the same mutation in *lmo2486*, despite all mutants originating from independent cultures derived from separate colonies. The probability of this occurring if the mutations were random is exceedingly low, suggesting a biological explanation. Indeed, the mutation is in a homopolymeric tract poly(A) of 8 nucleotides, which are known to have a substantially higher mutation frequency than expected by random chance (Gogol et al., 2007; Marvig et al., 2013; Orsi et al., 2010). The homopolymeric tracts poly(A) and poly(T) are overrepresented in many bacterial genomes and often at the start of genes. This is speculated to enable faster or a higher degree of adaptation in bacterial populations by gene inactivation from indels in such hypermutable sites (Orsi et al., 2010). Furthermore, internalin (encoded by the *inlA* gene), which is an important virulence factor in *L. monocytogenes* have been found to switch between an inactive frameshifted variant “A_6_” and a functional variant “A_7_” by addition/deletion of an adenine in a stretch of 7 adenine nucleotides (Orsi et al., 2010).

We also noted that the mutants were isolated from growth media containing 500 nM (3 isolates) and 1000 nM (1 isolate), corresponding to 4X and 8X the MIC_90_, displayed reduced resistance after being recultured in media in the absence of GarKS. After reculturing, the MIC_90_ was only 2-fold higher than that of the wild type, although the strains still harbored a mutated Lmo2486. This could suggest that transient gene regulation is an additional important factor.

We therefore used transcriptomics to investigate potential pathways to GarKS resistance development. RNA sequencing of this strain exposed to GarKS compared to unexposed controls revealed an upregulation of 23 genes. Among the most notable genes were members of a system resembling the phage shock protein response. Sequencing of spontaneous mutants resistant to GarKS identified mutations in a gene encoding the Psp protein Lmo2486, and its role in GarKS susceptibility was confirmed by cloning, protein expression, and susceptibility assays. As such, Lmo2486 alone is sufficient to explain the differences in GarKS susceptibility in the mutants. It is also interesting to note that mutations in a PspC-domain-containing protein have been reported to be associated with decreased GarKS susceptibility in *Lactococcus lactis* (Ovchinnikov et al., 2020).

The function of the Psp system in *L. monocytogenes* is poorly understood, and it is unclear how a disruption of Lmo2486 results in decreased GarKS susceptibility. The Psp proteins are thought to constitute a cell envelope stress response system, first described in *E. coli* by Brisette et al. (1990). They observed that infection by a filamentous phage f1 resulted in a large amount of a protein, PspA, being produced in response to the phage protein pIV, a pore-forming protein acting on the outer membrane (Brissette et al., 1990). Upregulation of PspA was also observed with environmental stress like heat, ethanol, and osmotic shock (Brissette et al., 1990). PspA is encoded by the *pspABCDE* operon, where PspBC likely detects an inducing signal that triggers the dissociation of a PspF-PspA complex and subsequent recruitment of liberated PspA to the inner membrane (Flores-Kim and Darwin, 2016). Recruitment of PspA to the inner membrane is believed to restore a compromised membrane and help reestablish the proton motive force by an unknown mechanism (Flores-Kim and Darwin, 2016).

Many variations to the Psp response have been found in bacteria, and these systems are poorly characterized in Bacillota (Ravi et al., 2024). *B. subtilis* encodes two separate and distinct Psp-like response systems, LiaIHGFSR and YvlABCD. In the former system, LiaRS is a typical two-component regulatory system (TCS) where LiaF is an inhibitor of LiaS under non-inducing conditions (Manganelli and Gennaro, 2017). Exposure to membrane-perturbing compounds such as daptomycin and nisin removes the inhibitory action of LiaF, which results in an increased expression of LiaIH. LiaH is a PspA ortholog and is recruited to the membrane by LiaI upon activation to restore membrane integrity. On the other hand, YvlABCD represents a second, less well-characterized, Psp-like system that is more directly homologous to the Psp system in *E. coli*. In contrast to the Lia system, expression of *yvl* is independent of LiaRS but is at least partly regulated by an extracytoplasmic function σ-factor (σ^W^), which is activated by cell envelope stress and cell wall synthesis inhibitors (Helmann, 2016). In this operon, YvlC is a PspC homolog, and YvlD is a phage holin-like protein, while YvlB contains a so-called “toastrack” (DUF4097) domain found in many Psp systems either alone or fused to a PspC domain (Ravi et al., 2024). The toastrack domain has been hypothesized to function as a protein-protein interaction platform, either in the cytosol or anchored to the intracellular side of the membrane (Ravi et al., 2024). Similarly to YvlABCD, the *lmo2487-6-5-4* operon of *L. monocytogenes* encodes a holin-like protein (Lmo2484), a PspC-domain-containing protein Lmo2485, and two toastrack-domain-containing proteins, Lmo2486 and Lmo2487, in which the former is also fused to a PspC domain. Most importantly, Lmo2486 was directly linked to GarKS resistance. The organization of the *psp* operons in the three species is shown in Figure 2.

**Figure 2.**
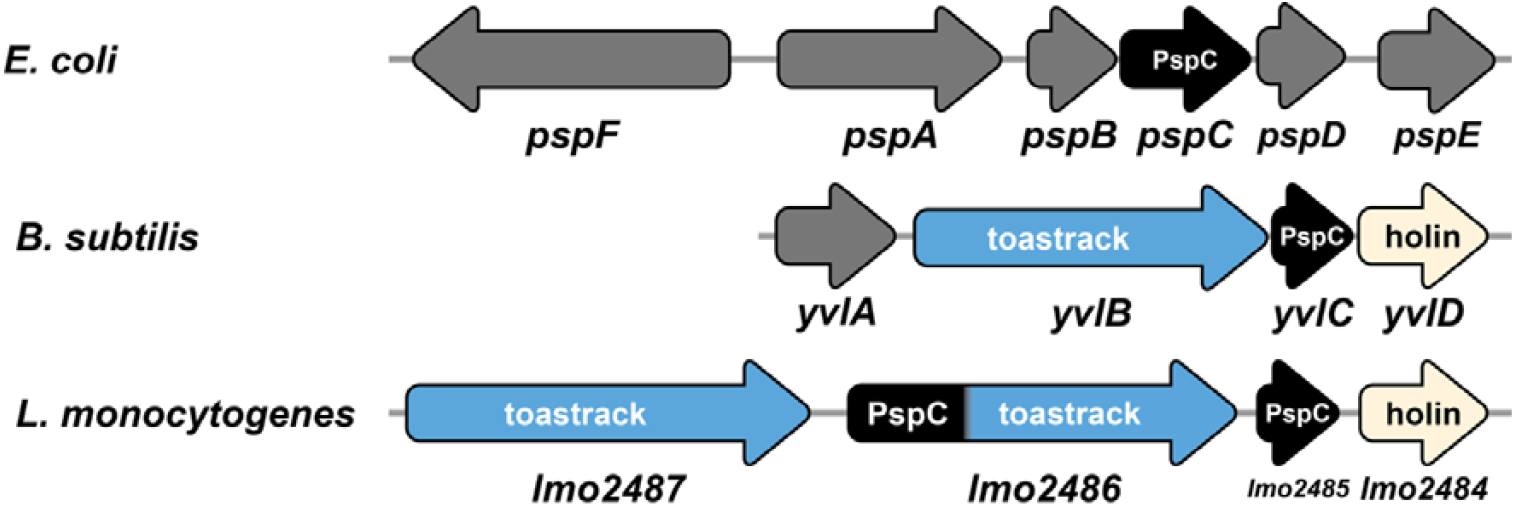
Operon organization of *psp* in *E. coli* (strain K-12, GenBank accession U00096), *B. subtilis* 168 (GenBank AL009126) and *L. monocytogenes* EGD-e and RO15. Labels and colors inside arrows indicate the characteristic conserved domain encoded by each gene: toastrack (DUF4097) in blue; phage shock protein C (Phageshock_PspC_N) in black; mycobacterial 4 TMS phage holin, superfamily IV in beige/cream; and unknown/other in dark gray.

Although the function of the Yvl proteins is not known, YvlBC could be a stress sensor and regulator like PspBC. Analogously, Lmo2487 (toastrack) and Lmo2485 (PspC domain) could serve a similar function as PspBC/YvlBC in *L. monocytogenes*. However, in this organism the *psp* operon does not encode any putative inhibitors or negative regulators, which suggests that *L. monocytogenes* has a divergent Psp system that functions differently or relies on other effectors. Although speculative, Lmo2486 could serve this function either by direct interaction with an Lmo2487-Lmo2485 complex or by a competitive interaction with the inducing signal. A disruption mutant of *lmo2486* would be expected to have a higher basal activation of the Psp system, which would then confer a protective effect by an unknown mechanism.

Similar to *B. subtilis*, a LiaFSR-orthologous TCS is found in *L. monocytogenes* (Lmo1020-1022), which is also associated with cell envelope stress (Fritsch et al., 2011). Furthermore, genes known to be regulated by this TCS are *lmo0954-0955, lmo1966-1967, lmo2487–2484, lmo2567*, and *lmo2568* (Nielsen et al., 2012). Many of which were found to be upregulated by GarKS exposure in this study (Figure 1, Table 4), suggesting that the LiaRS TCS is activated by GarKS. The highest fold-change difference upon GarKS exposure was observed for a LiaSR-regulated gene, *lmo2568*. This gene encodes an YxeA family protein known to be upregulated by cationic peptides in *Bacillus* and is part of a regulon involved in cell wall modifications in response to antimicrobials active on the cell envelope (Joseph et al., 2004; Pietiäinen et al., 2005; Wecke et al., 2006). The putative operon *lmo1966-lmo1967* encodes proteins similar to XpaC in *Bacillus subtilis* and TelA/YaaN involved in tellurite resistance, respectively. However, the function of these proteins in *Listeria* has not been well established. The screening of a *mariner* transposon library of *L. monocytogenes* EGDe found that a disruption mutant of *lmo1967* had a 4-fold increased susceptibility to nisin and a 2-fold increase for tellurite (Collins et al., 2010). The Lmo1967 homolog in *B. subtilis* (YaaN) is also upregulated by the cationic antimicrobial peptide poly-L-lysine (Pietiäinen et al., 2005). It seems likely that transient upregulation of Lmo1967 in *L. monocytogenes* can contribute to GarKS tolerance in wild-type cells, possibly by active export of toxic charged molecules out of the cell. However, the mechanism of Lmo1967 in resistance to these compounds is not known.

In conclusion, *L. monocytogenes* isolates from food, fish, and fish processing environments are highly susceptible to the bacteriocin GarKS. Furthermore, resistance to GarKS is infrequent, and only an 8-fold decrease in susceptibility was seen in our study. The phage shock protein response of *L. monocytogenes* is directly involved in GarKS susceptibility, and in particular Lmo2486, a PspC-domain-containing protein. Lmo2486 and other members of this stress response system may serve as novel antimicrobial targets. GarKS is a highly promising antimicrobial for use in food preservation, particularly in combination with other food processing technologies as part of a hurdle strategy to further reduce or eliminate spoilage bacteria and foodborne pathogens (e.g., from perishable food products such as fresh fish and cold-smoked salmon).

## Declarations

### Ethical approval

Not applicable.

### Data availability

The sequencing data generated in this research is available from the European Nucleotide Archive (ENA) under Project accession no. PRJEB93936.

